# Brainstem control of urethral sphincter relaxation and scent marking behavior

**DOI:** 10.1101/270801

**Authors:** Jason Keller, Jingyi Chen, Sierra Simpson, Eric Hou-Jen Wang, Varoth Lilascharoen, Olivier George, Byung Kook Lim, Lisa Stowers

## Abstract

Urination may occur either reflexively in response to a full bladder or deliberately irrespective of immediate need. Voluntary control is desired because it ensures that waste is expelled when consciously desired and socially appropriate^1,2^. Urine release requires two primary components: bladder pressure and urethral relaxation^1–3^. Although the bladder contracts during urination, its slow smooth muscle is not under direct voluntary control and its contraction alone is not sufficient for voiding. The decisive action of urination is at the urethral sphincter, where striated muscle permits fast control. This sphincter is normally constricted, but relaxes to enable urine flow. Barrington’s nucleus (Bar, or pontine micturition center) in the brainstem is known to be essential for the switch from urine storage to elimination^4–7^, and a subset of Bar neurons expressing corticotropin releasing hormone (Bar^CRH^) have recently been shown to promote bladder contraction^8–10^. However, Bar neurons that relax the urethral sphincter to enable urination behavior have not been identified. Here we describe novel brainstem neurons that control the external urethral sphincter. We find that scent marking behavior in male mice depends upon a subpopulation of spatially clustered Bar neurons that express high levels of estrogen receptor 1 (Bar^ESR1^). These neurons are glutamatergic, project to urinary nuclei in the spinal cord with a bias towards sphincter-inhibiting interneurons, and their activity correlates with natural urination. Optogenetic stimulation of Bar^ESR1^ neurons rapidly initiates sphincter bursting and efficient voiding in absence of sensory cues in anesthetized and behaving animals. Conversely, inhibiting the activity of these neurons prevents olfactory cues from promoting scent marking behavior. The identification of Bar^ESR1^ cells provides an expanded model for the supraspinal control of urination and its dysfunction.

---

Supraspinal neurons that relax the urinary sphincter to facilitate urine flow (Figure 1a) have not been identified. Bar contains at least three different cell types defined by physiology^11–14^, gene expression^10,15^, and histology^15–17^. Among these, Bar^CRH^ are best-studied and increase their firing rate during anesthetized bladder and colon distension^3^ as well as awake, diuretic-induced urination. Moreover, they increase bladder pressure following optogenetic stimulation^10^. However, our initial tests and a previous study of Bar^CRH^ neural function^10^ showed modest effects on urination in awake animals. About half of the Bar neurons projecting to the spinal cord lack CRH expression^9^, but their molecular identity and function is unknown^15^. We took a candidate approach to identify molecular markers for Bar neurons that may function to promote urinary sphincter relaxation, and focused on estrogen receptor 1 (ESR1, ESRα), as it is expressed in a subset of Bar cells in both mice^17^ and primates^18^. It is unknown if ESR1 marks a cell type distinct from Bar^CRH^. Immunostaining with αESR1 in CRH-Cre x ROSA-LSL-tdTomato (CRH-tdT) individuals confirmed a small population (~200 cells) expressing high amounts of ESR1 (Bar^ESR1^ neurons, Fig. 1b-e). The majority of Bar^ESR1^ neurons (~3/4 of the Bar^ESR1^ population) do not overlap with CRH-tdT, and the overlapping minority likely represents an upper bound on co-expression since tdT integrates CRH promoter activity over the lifetime of the animal (Fig. 1e). Bar^ESR1^ neurons are found in a dorsal cluster within the Nissl-defined ovoid Bar nucleus, whereas Bar^CRH^ neurons are more numerous (~500 cells^10^), ventrally biased, and extend further along the rostrocaudal axis beyond traditional, Nissl-defined Bar borders^19^ (Fig. 1c-d). Moreover, in ESR1-Cre mice^20^, 96.8 % of Bar^ESR1^ neurons (N=3 mice) overlap with reporter expression (Extended Data 1a), confirming that the CRH and ESR1 promoters are active in largely independent Bar populations.

**Fig 1.**
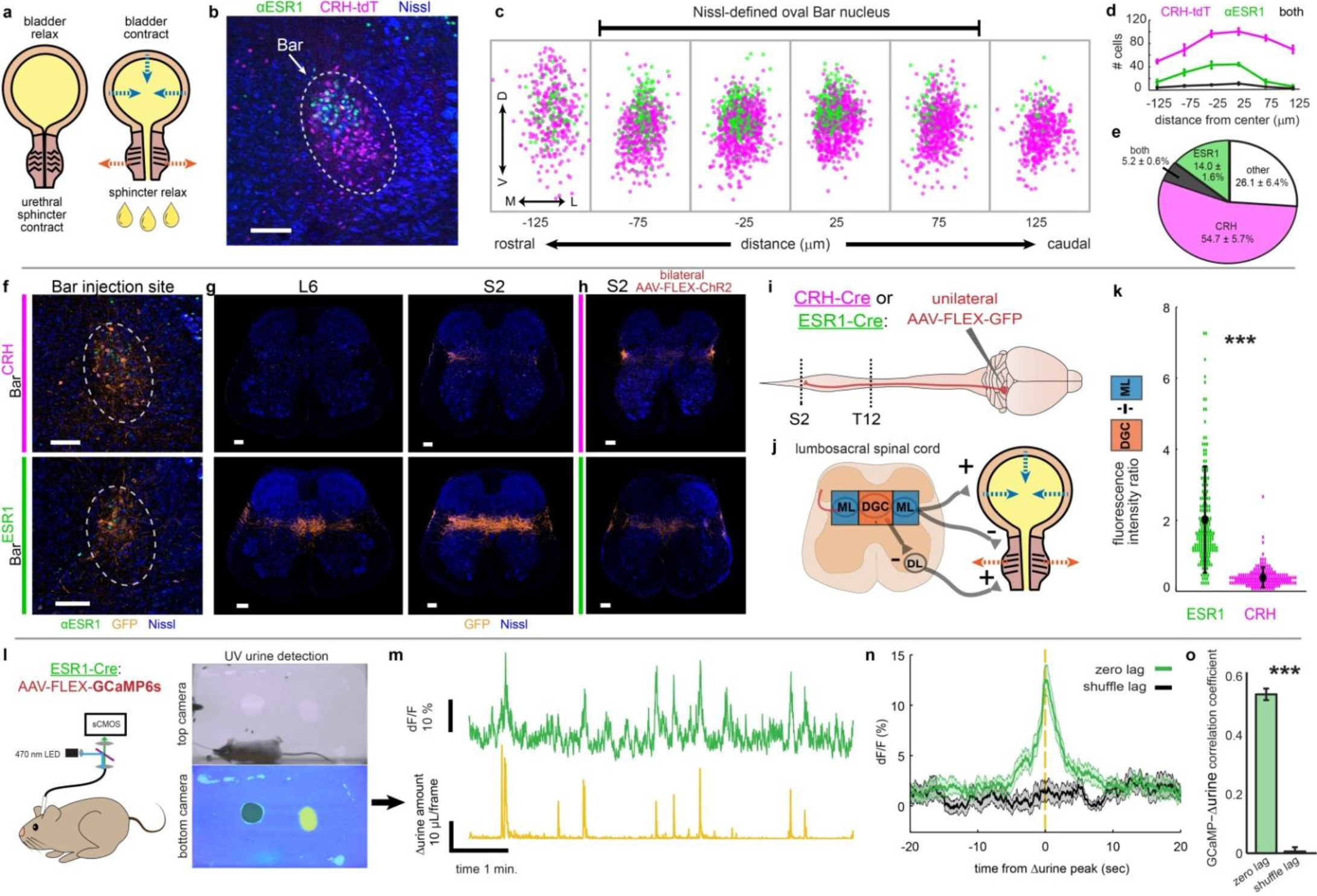
A novel cell type in Barrington’s nucleus displays features indicating a role in urination. **a,** Urination requires sphincter relaxation. **b,** ESR1-immunostaining in Bar (dotted oval) in CRH-tdT mice. **c,** Rostrocaudal overlay of αESR1 cells (green) in Bar registered to centroid of CRH-tdT cells (magenta). **d,** Cell counts, and **e**, cell percentages in Bar (mean ± s.e.m., N = 6). **f**, GFP expression at Bar injection site in CRH-Cre (top) or ESR1-Cre (bottom) individuals. **g**, Axonal projections in lumbosacral spinal cord (right L6, left S2) for injections in (f). **h**, Axonal projections in lumbosacral S2 spinal cord for injection sites in Fig. 2b. **i**, Schematic for identifying Bar cell type axonal projections to spinal cord. **j**, Simplified urinary circuitry in the lumbosacral spinal cord. ML = mediolateral column, DGC = dorsal grey commissure, DL = dorsolateral nucleus. **k**, Quantification of Bar^ESR1^ and Bar^CRH^ axonal projections in lumbosacral spinal cord. Points are individual sections, thick black line is mean ± s.e.m for Bar^CRH^ (magenta, N = 10), Bar^ESR1^ (green, N = 10). **l**, Schematic of fiber photometry experiment and example urine quantification with control odor (black shading) and female odor (yellow shading) on bottom camera view. **m**, Example Bar^ESR1-GCaMP6s^ fluorescence (top) and derivative of urine detection (Δurine, bottom). **n**, GCaMP6s fluorescence synchronized to Δurine peaks (green) or at shuffled times (black) for all mice (mean ± s.e.m, N = 73 urination events from 10 mice). **o**, Correlation coefficient between GCaMP6s and Δurine traces at zero lag (green) and random lag (black) for all mice (mean ± s.e.m., same events as panel n). Scale bars = 100 μm. ***p<0.001 (Wilcoxon rank sum).

To investigate the potential for Bar^ESR1^ neurons to relax the urethral sphincter, we evaluated their neurotransmitter identity and anatomical connections to the lower urinary tract. Immunostaining for αESR1 in Vgat-Cre and Vglut2-Cre mouse lines crossed to fluorescent reporters, as well as in-situ hybridization, revealed that the majority of Bar^ESR1^ neurons express Vglut2 (93.6 % reporter overlap, N = 3 mice) and not Vgat (2.2 % reporter overlap, N = 4 mice; Extended Data 1a-k). Injection of the retrograde tracer CTB into the lumbosacral spinal cord resulted in co-expression with Bar^ESR1^ cells, indicating their direct projections to urinary targets (Extended Data 1l-n). To further investigate Bar^ESR1^ axonal projections, we unilaterally injected AAV expressing Cre-dependent GFP into the Bar of ESR1-Cre or CRH-Cre animals, and imaged the lower thoracic to sacral spinal cord (Fig. 1f,g,i). The lumbosacral mediolateral column (ML) contains preganglionic autonomic neurons that excite the bladder (along with intermingled interneurons)^1,21,22^, and the lumbosacral dorsal grey commissure (DGC) contains interneurons that directly inhibit (relax) sphincter motorneurons of the dorsolateral nucleus via Bar input^21,23^ (Fig. 1j). Consistent with the known role in bladder pressure regulation, Bar^CRH-GFP^ axons showed a dense focal projection to the ML (Fig. 1f-g, k) with only sparse fibers arching further medially or to thoracolumbar levels T13-L2 (Extended Data 2a-b). Bar^ESR1-GFP^ axons projected similarly across the lumbosacral ML, with additional lighter fibers seen in the thoracolumbar ML (Extended Data 2c). However, they also provided much denser innervation of the sphincter-inhibiting DGC, extending rostrally from the proposed L3-L4 burst generator^24^ to mid-sacral levels (Fig. 1f-g,k, Extended Data 2c). Bilateral labeling of Bar^ESR1^ or Bar^CRH^ neurons with a second Cre-dependent virus (AAV-FLEX-ChR2) confirmed the same projection patterns (Fig.1h, Extended Data 2b,c). Thus, the cell body distribution, molecular expression, and efferents of Bar^ESR1^ neurons indicate that they constitute an uncharacterized cell type within Bar^15^, distinct from Bar^CRH^ neurons.

To determine the temporal activation of Bar^ESR1^ cells in relationship to natural urination behavior, we unilaterally injected Bar with AAV-FLEX-GCaMP6s in ESR1-Cre animals and imaged population calcium activity with fiber photometry (Fig.1l). Our behavioral assay enabled freely moving scent marking behavior while quantifying the timing and abundance of urination events (Extended Data 3). We found activity in Bar^ESR1^ cells to be highly correlated with detected urination events, compared to randomly chosen intervals (Fig 1m-o). Altogether, we find that ESR1 defines a novel cell type in Bar that displays features consistent with a role in urination.

Bar^CRH-ChR2^ photostimulation was previously shown to drive bladder pressure increases during urethane-anesthetized cystometry^10^, but the sufficiency of these cells in awake urination has not been characterized. To determine if either of these distinct Bar populations promote urination in behaving animals, we first bilaterally infected Bar^ESR1^ or Bar^CRH^ neurons with AAV-FLEX-ChR2 or -GFP (Bar^ESR1-ChR2^, Bar^ESR1-GFP^, or Bar^CRH-ChR2^, Fig. 2a-c) and performed slice recordings to confirm that both Bar^ESR1-ChR2^ and Bar^CRH-ChR2^ neurons reliably responded to photostimulation at frequencies previously used in electrical stimulation (Extended Data 4). We then quantified and compared the rate and amount of urine induced by photostimulation in awake, freely-moving individuals. While photostimulation of GFP-infected individuals produced no effect on urine excretion, Bar^ESR1-ChR2^ stimulation led to robust, frequency-dependent urine volume released, following light onset with a mean latency of 2.1 seconds (Fig. 2d-h; Suppl. Video 1). Over 96% of Bar^ESR1-ChR2^ stimulation trials at 10-50 Hz resulted in urination (Fig. 2d,f). In comparison, photostimulation of Bar^CRH-ChR2^ neurons during freely-moving behavior had a much smaller effect on urination despite generally higher ChR2 viral infection levels (Fig. 2c-h; Suppl. Video 2). Less than 37% of Bar^CRH-ChR2^ stimulation trials at 10-50 Hz resulted in the voiding of urine (Fig. 2f). Of this subset, the latency and amount of urine produced differed from Bar^ESR1-ChR2^ at all frequencies tested (Fig.2d-h). We additionally investigated the extent to which Bar^ESR1^ and Bar^CRH^ neural activity could initiate voiding without conscious sensory input. Photostimulation under isoflurane anesthesia, known to depress reflex urination^25–27^, resulted in urine voiding in 43% of the Bar^ESR1-ChR2^ trials, but only 6% of the Bar^CRH-ChR2^ trials, with none of the Bar^CRH-ChR2^ voids occurring during the photostimulus window (Fig. 2i; Suppl. Video 3). This indicates that Bar^ESR1^ neuronal activity is sufficient to trigger rapid and efficient urination and hints at a distinct mechanism from neighboring Bar^CRH^ activity that is known to increase bladder pressure.

**Fig 2.**
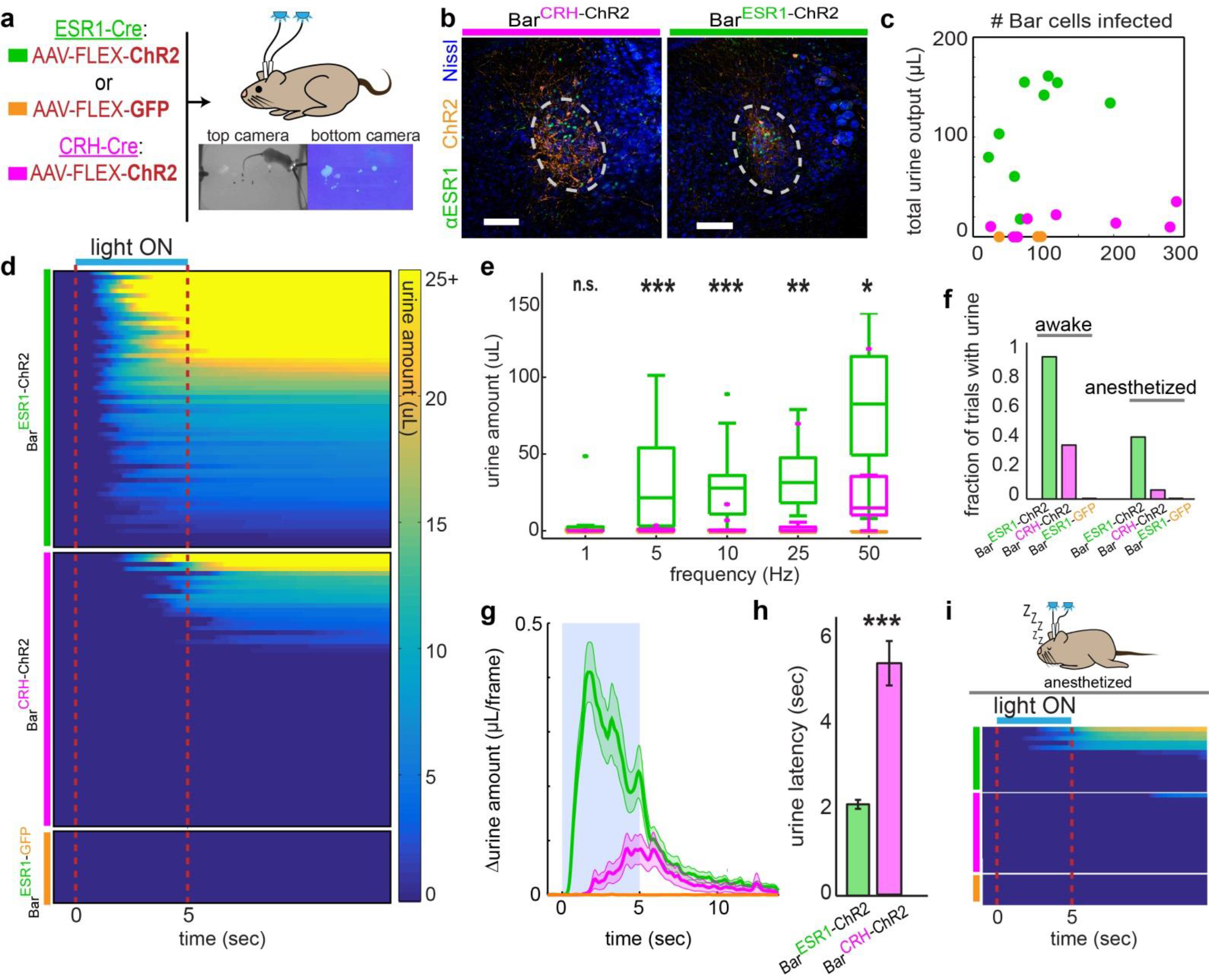
Photostimulation of Bar^ESR1^ neurons induces efficient urination in awake and anesthetized animals. **a,** Schematic of optogenetic stimulation and example urine detection. **b,** Example ChR2 expression in CRH-Cre (left) or ESR1-Cre (right) individuals. **c,** Total urine output across all trials for each individual versus ChR2 expression. **d,** Heatmap of urine output following awake photostimulation for all trials >10Hz (N = 10 Bar^ESR1-ChR2^, 10 Bar^CRH-ChR2^, 3 Bar^ESR1-GFP^ mice), sorted by decreasing total urine amount. **e,** Urine amounts at different photostimulation frequencies: boxplots show median, 25^th^/75^th^ quartiles, ranges, and outliers (same mice as panel d). **f,** Fraction of trials with photostimulated urine detected in panels (d), awake, and (i), anesthetized. **g,** Δurine amount around photostimulation (blue shading; same mice as panel d). **h,** Urination latency after photostimulation (N = 10 Bar^ESR1-ChR2^, 10 Bar^CRH-ChR2^ mice). **i,** Heatmap of urine output around anesthetized photostimulation for all trials (N = 7 Bar^ESR1-ChR2^, 8 Bar^CRH-ChR2^, 3 Bar^ESR1-GFP^ mice). Scale bars = 100 μm. *p<0.05, **p<0.01, ***p<0.001, n.s. p>0.05 (Wilcoxon rank sum for Bar^ESR1-ChR2^ compared to Bar^CRH-ChR2^). Colors for all panels: green = Bar^ESR1-ChR2^, magenta = Bar^CRH-ChR2^, orange = Bar^ESR1-GFP^.

To directly test the effect of Bar^ESR1^ and Bar^CRH^ neurons on urinary muscle targets, we performed urethral sphincter electromyography (EMG) and cystometry (bladder filling and pressure recording) under isoflurane anesthesia (Fig 3a). We perfused saline at a constant rate into the bladder to stimulate reflex voiding and observed natural cycles of bladder pressure increase and associated external urethral sphincter bursting muscle patterns, which correlated with voiding and subsequent bladder pressure decrease (Fig.3b). In rodents, these bursting contractions interspersed with periods of muscle relaxation are believed to enable efficient urine flow through the narrow rodent urethra^28–30,1^. Following observation of regular cystometry cycles, we stopped the saline pump when the bladder was filled to 75% of the volume observed to trigger reflex urination, and initiated 5 seconds of photostimulation (Fig. 3b, blue arrows). We found that both Bar^ESR1-ChR2^ and Bar^CRH-ChR2^ photostimulation produced reliable, time-locked bladder pressure increases at similar latencies (Fig. 3c-d, Extended Data 5a). The initial latency and slope of the bladder pressure increase by stimulation of each cell type was indistinguishable by our analysis; however, the peak pressure and end pressure (25 seconds after stimulus onset) were significantly less for Bar^ESR1-ChR2^ photostimulation, which reflects abundant urine release (Fig. 3c-f, Extended Data 5a). This increase in urination coincided with a reliable bursting pattern of sphincter activity only during Bar^ESR1-ChR2^ photostimulation (Fig. 3b,g-k; Extended Data 5e-f, 6b). Using wireless pressure recording from the corpus spongiosum as a proxy for the urethral pressure, we also found similar urethral sphincter bursting patterns to occur during the natural behavior response to odor cues (Extended Data 6a-d). Notably, the spectral power seen during the Bar^ESR1-ChR2^ photostimulation bursts mimicked wirelessly recorded pressure during scent-marking urination (Extended data 6c,d,f). During the photostimulated sphincter bursts we observed pulsatile urination (Suppl. Video 4) which was dependent on bladder fill level (Fig. 3g-h, Extended Data 5b). Frequency analysis of the sphincter EMG signal shows that 85% of the Bar^ESR1-ChR2^ stimulations resulted in sphincter relaxation/bursting and associated voiding (Fig.3g-k; Extended Data 5a,b). Additionally, we observed burst-like EMG responses in the absence of bladder contractions during Bar^ESR1-ChR2^ photostimulation on a subset of trials when the bladder was only filled to 10% of reflex urination volume (Extended Data 5c). In contrast, photostimulation of Bar^CRH-ChR2^ neurons produced either no detectible change in sphincter activity, tonic sphincter discharge (constriction), or rare irregular bursting (13% of trials), which was always preceded by tonic activity (Fig. 3b,g-k; Extended Data 5b). This tonic activity increase was characteristic of a spinal guarding reflex mediated through bladder afferents to prevent urination during bladder distension. When Bar^CRH-ChR2^ photostimulation ceased, the bladder usually returned to the same pressure level observed prior to Bar^CRH-ChR2^ activity (Fig 3c-d,f), indicating significant urine was not normally released (Suppl. Video 5). Overall, these results indicate that activity from both Bar populations equally increase bladder pressure, but only Bar^ESR1^ neurons actively promote bursting of the urethral sphincter muscle to enable efficient urine flow.

**Fig 3.**
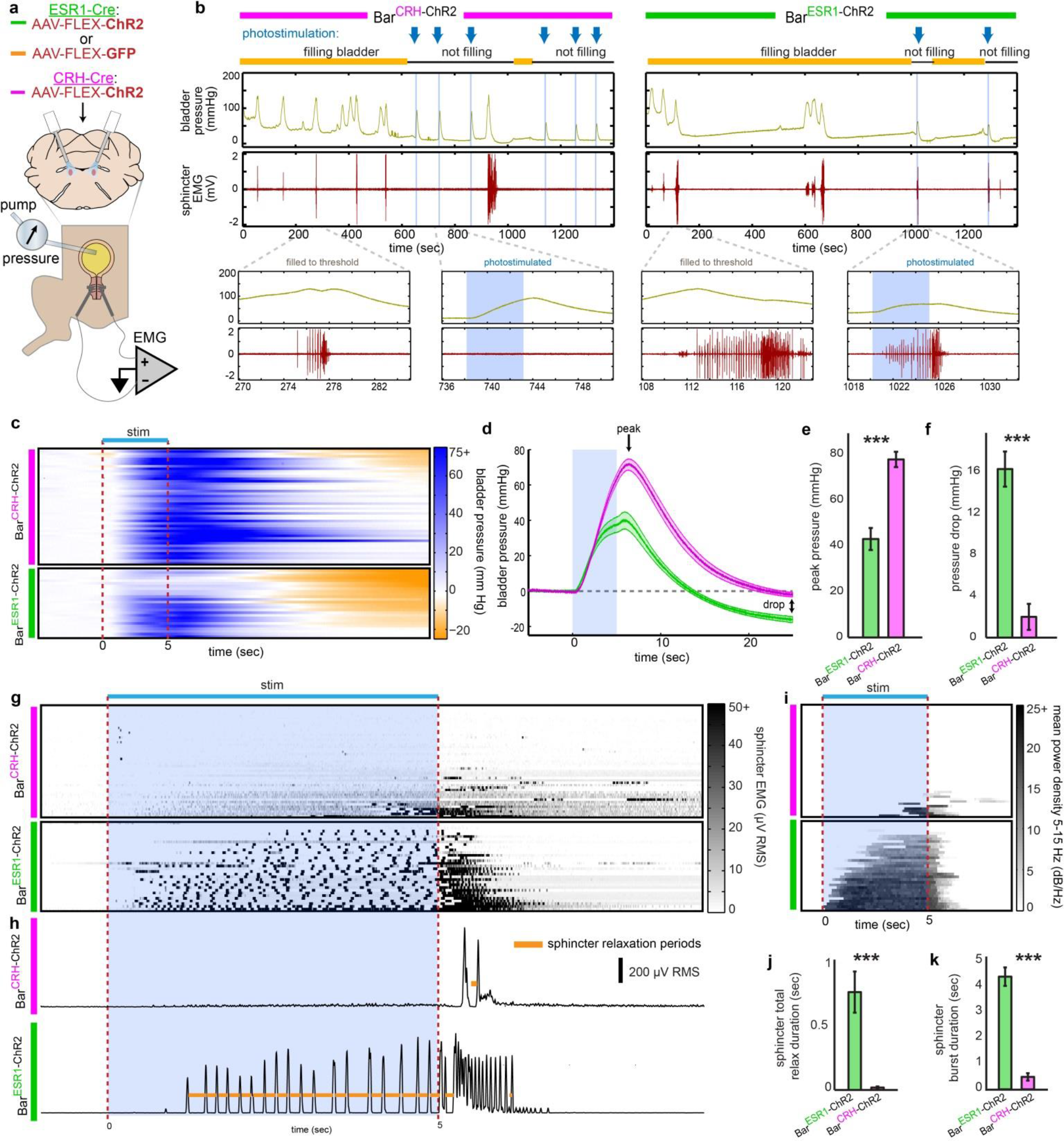
Bar^ESR1^ neurons control the urethral sphincter. **a,** Schematic for optogenetic Bar stimulation during cystometry. **b,** Representative raw bladder pressure and sphincter EMG traces for Bar^CRH-ChR2^ (left) and Bar^ESR1-ChR2^ (right) individuals. Blue arrows and shading indicate photostimulation times, and yellow/black lines denote cystometry pump on/off. Top traces are 20 minutes; bottom traces show 15 second detail when the bladder is filled to threshold (no photostimulation) versus when Bar is photostimulated. **c,** Heatmap of bladder pressure around photostimulation for all trials (N = 5 Bar^CRH-ChR2^, top, and 3 Bar^ESR1-ChR2^, bottom, mice) in order of decreasing maximum pressure drop. **d,** Bladder pressure for data in panel (c), showing peak and final pressure (mean ± s.e.m., green = Bar^ESR1−ChR2^, magenta = Bar^CRH-ChR2^). **e,** Peak bladder pressure from (d), mean ± s.e.m. **f,** Bladder pressure drop from (d), mean ± s.e.m. **g,** Heatmap of sphincter RMS EMG around photostimulation for all trials (N = 5 Bar^CRH-ChR2^, top, and 6 Bar^ESR1-ChR2^, bottom, mice) in increasing mean voltage order. **h,** Example RMS EMG traces from a single trial in (g), showing calculated sphincter relaxation periods (orange) between bursts. **i,** Heatmap of mean EMG power density at bursting frequencies (5-15 Hz) for trials in (g) (Bar^CRH-ChR2^, top, and Bar^ESR1-ChR2^, bottom). **j,** Total sphincter relaxation time for trials in (g) (mean ± s.e.m). **k,** Sphincter burst duration for trials in (i) (mean ± s.e.m). ***p<0.001 (Wilcoxon rank sum).

Upon detecting the odor of a female, males promptly urinate to show their command of the territory and advertise their availability to mate^31–33^. We established a rapid behavioral assay that compares the voluntary baseline urination rate (two minutes in the presence of a control odor) to the rate during the subsequent two minutes, in the presence of motivating female urine odor (Figure 4a-c, Suppl. Video 6). The reliable and rapid change in the amount of urine marks in response to female urine indicates that olfactory cues access circuits which relax the urethral sphincter and generate voluntary urination. To test if Bar neurons are necessary for this response, we bilaterally infected them with AAV-FLEX-hM4Di in ESR1-Cre or CRH-Cre mice (Bar^ESR1-hM4Di^ or Bar^CRH-hM4Di^; Fig. 4d, Extended Data 7a,d). Individuals were then injected with either CNO or saline on alternate days and assayed for their urination rate in the presence of female urine. Female-odor evoked urination was reversibly diminished following CNO injections in Bar^ESR1-hM4Di^ but not Bar^CRH-hM4Di^ or wild-type control mice (Fig. 4e-f), again despite higher viral infection levels in CRH-Cre mice (Extended Data 7d), and without affecting locomotion or odor sampling (Extended Data 7b-c). A previous study found a subtle effect on urination from Bar^CRH-hM4Di^ inhibition at a much longer 2-hour timescale, which we replicated here (Extended Data 7e) and suggests a modulatory role for Bar^CRH^ neurons. We additionally assayed for necessity of Bar^ESR1^ neurons at faster timescales by bilaterally injecting them with AAV-FLEX-ArchT (Bar^ESR1-ArchT^ mice; Fig. 4g, Extended Data 8a). We compared urination during 2 minutes of photoinhibition with female odor present to an additional 2 minutes immediately after without photoinhibition (Fig. 4h). Sniffing of the female odor did not differ during and after photoinhibition, but urination was largely inhibited during the photoinhibition window (Fig. 4 h-k, Extended Data 8b, Suppl. Video 7). Most trials with female odor, but not with control odor, resulted in urination within seconds of light termination. This suggests that the immediate urine release resulted from priming by odor cues rather than trivial rebound activity upon the cessation of photoinhibition (Fig.4h-i; Extended Data 8c). Finally, photoinhibition during cystometry revealed that ongoing Bar^ESR1^ activity is necessary to maintain sphincter bursting (Extended Data 8d-e). Together, our experiments indicate that Bar^ESR1^ neurons are essential for urethral inhibition and voluntary urination promoted by olfactory cues in male mice (Extended Data 9). Bar^ESR1^ neurons serve as a potential new target to study a variety of urinary disorders^1^ as well as a crucial node in a tractable motivated behavior with well-defined olfactory input and relatively simple muscle output.

**Fig 4.**
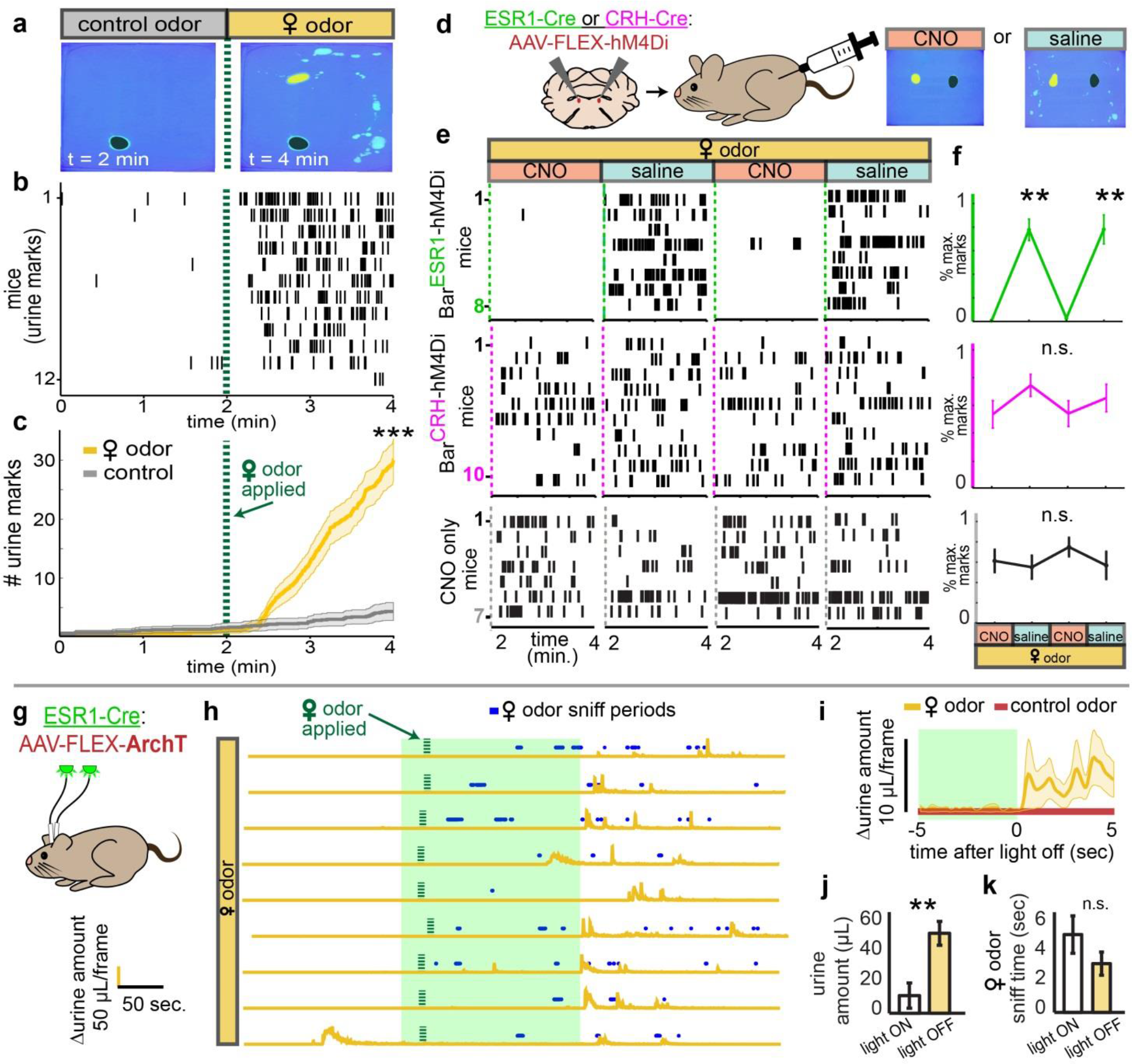
Bar^ESR1^ neurons are necessary for rapid, odor-evoked urination. **a,** Scent marking behavior in wild-type mice. Left: after 2 min. exposure to control odor (black shading), right: after additional 2 min. with female odor (yellow shading). **b,** Raster plot of urine marks detected. **c,** Urine marks during habituation with control odor only (grey) or with female odor (yellow) (mean ± s.e.m., N = 12 mice). ***p<0.001 (Wilcoxon signed rank) for number of urine marks at 2 min. and 4 min. **d,** Schematic of chemogenetic inhibition of Bar^ESR1^ during scent marking urination. **e,** Raster plots of urine marks on consecutive days with either CNO or saline (Bar^ESR1-hM4Di^, top, Bar^CRH-hM4Di^, middle, CNO-only control, bottom). **f,** Percentage of maximum urine marks across all CNO or saline days for, top, Bar^ESR1-hM4Di^ (N = 8), middle, Bar^ESR1-hM4Di^ (N = 10) and, bottom, CNO control (N = 7) mice (mean ± s.e.m.). **p=0.01, n.s. p>0.05 (Friedman’s with Dunn-Sidak posthoc) for differences between saline and CNO days. **g,** Schematic of optogenetic inhibition of Bar^ESR1^ during scent marking urination. **h,** Δurine amount around 2 min. photoinhibition period. Female odor presented within 15 seconds of light on, and subsequent sniff periods shown in blue. N = 9 trials from 3 mice. **i,** Δurine amount ± 5 sec. from end of photoinhibition for control odor and female odor (mean ± s.e.m., N = 9 total trials from 3 mice). **j, k,** Urine amount (k) and female odor sniff time (l) during 2 min. photoinhibition period and 2 min. immediately following (mean ± s.e.m., N = 9 trials). **p<0.01, n.s. p>0.05 (Wilcoxon rank sum). Green shading denotes photoinhibition periods.

## Author contributions and acknowledgments

J.A.K., J.C., and L.S. designed the study, analyzed the data, and wrote the manuscript.S.S. aided in the cystometry (O.G. supported S.S.). E.H.W and B.K.L. aided in the fiber photometry. V.L. performed the slice physiology. All other experiments were performed by J.A.K. and J.C. J.A.K. was supported by NSF-GRFP grant DGE-1144086.

---

**Extended Data Fig 1.**
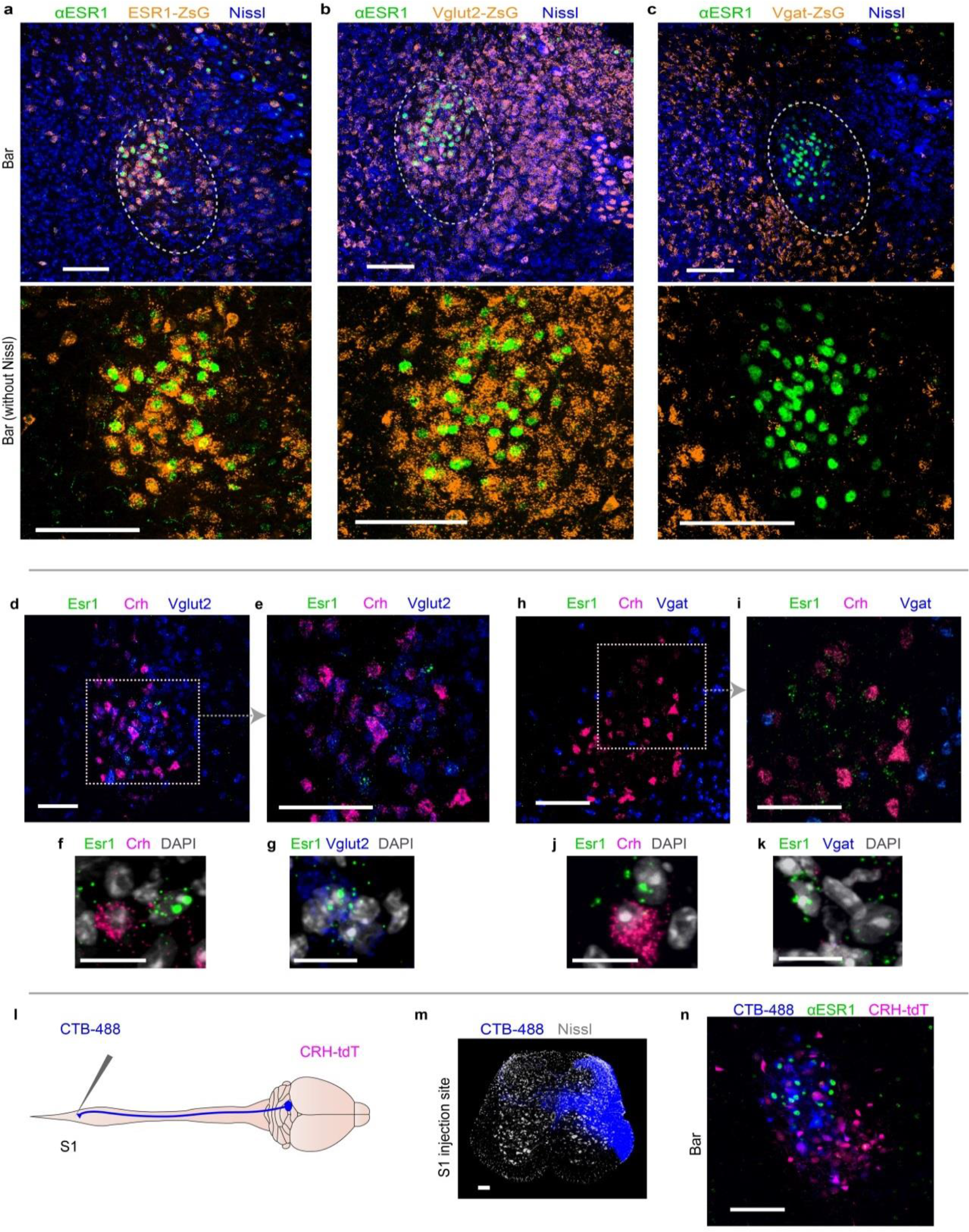
Neurotransmitter identity and direct spinal projections of Bar^ESR1^ neurons. **a,** αESR1 overlap with ESR1-ZsGreen (Ai6) genetic reporter. Bottom is larger view without Nissl. **b,** αESR1 overlap with Vglut2-ZsGreen genetic reporter. **c,** αESR1 overlap with Vgat-ZsGreen genetic reporter. **d,** RnaScope in-situ hybridization of Crh/Esr1/Vglut2 mRNA in Bar region of a wild-type male mouse, 20X objective. **e,** Larger view of dotted area in (d), 40X objective. **f, g,** Close-up views of individual cells in (e), with DAPI counterstain. **h-k,** same as (d)-(g), but with Vgat mRNA probe. **l,** Schematic of CTB injection into S1 spinal cord. **m,** CTB injection site. **n,** Retrograde CTB labeling in Bar with αESR1 and CRH-tdT. Dotted ovals delineate Bar. Scale bars = 100 μm, except panels f/g/j/k, scale bars = 20 μm.

**Extended Data Fig 2.**
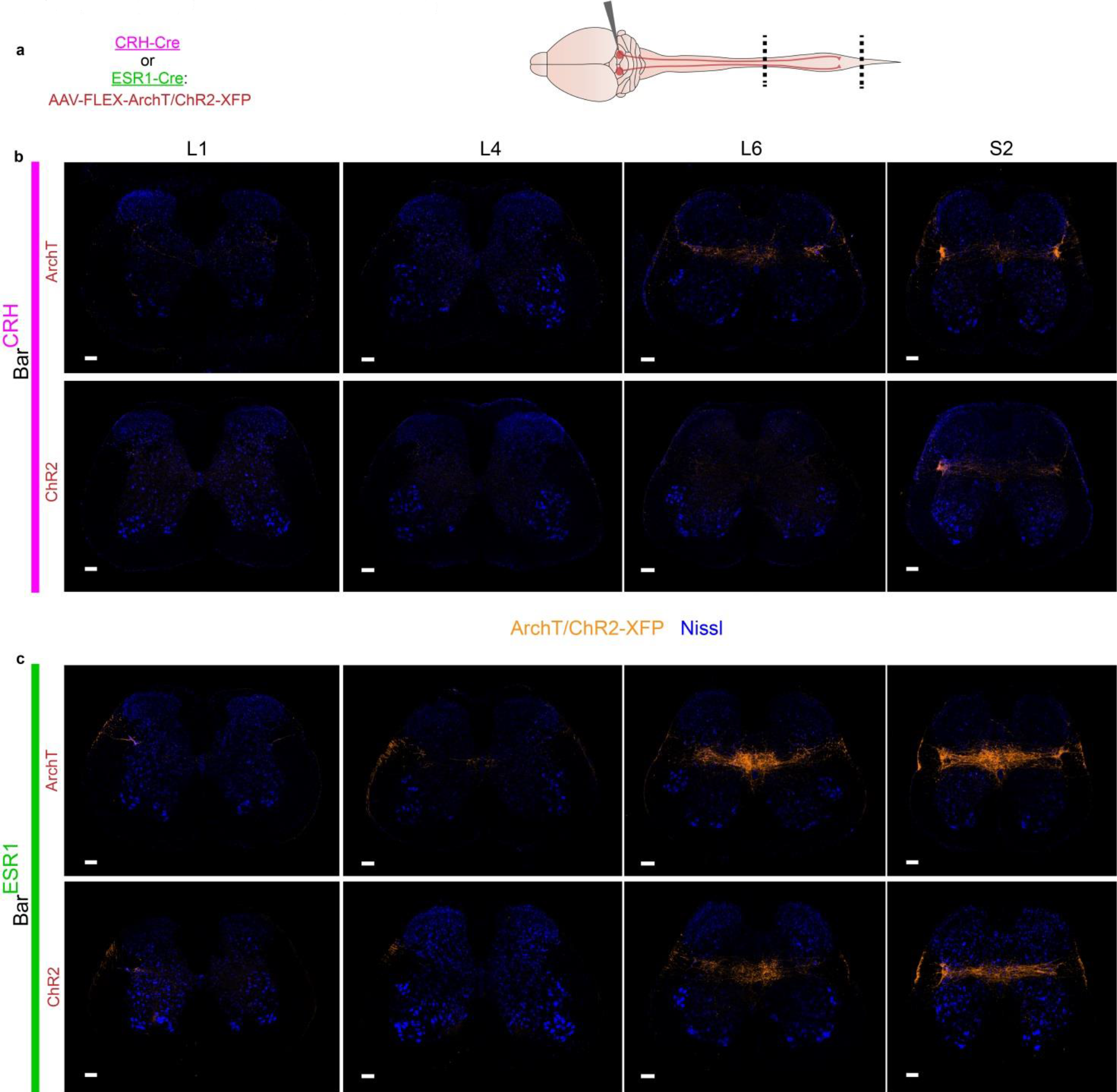
Bar^ESR1^ and Bar^CRH^ projections to urinary nuclei in the spinal cord. **a,** Schematic for testing Bar cell type axonal projections to spinal cord. **b,** Representative Bar^CRH^ axon projections at the L1/L4/L6/S2 spinal cord levels, separate from example in Fig. 2f-h. Top is ArchT virus, bottom is ChR2 virus. **c,**. Same as (b), but for Bar^ESR1^ axon projections. Scale bars = 100 μm.

**Extended Data Fig 3.**
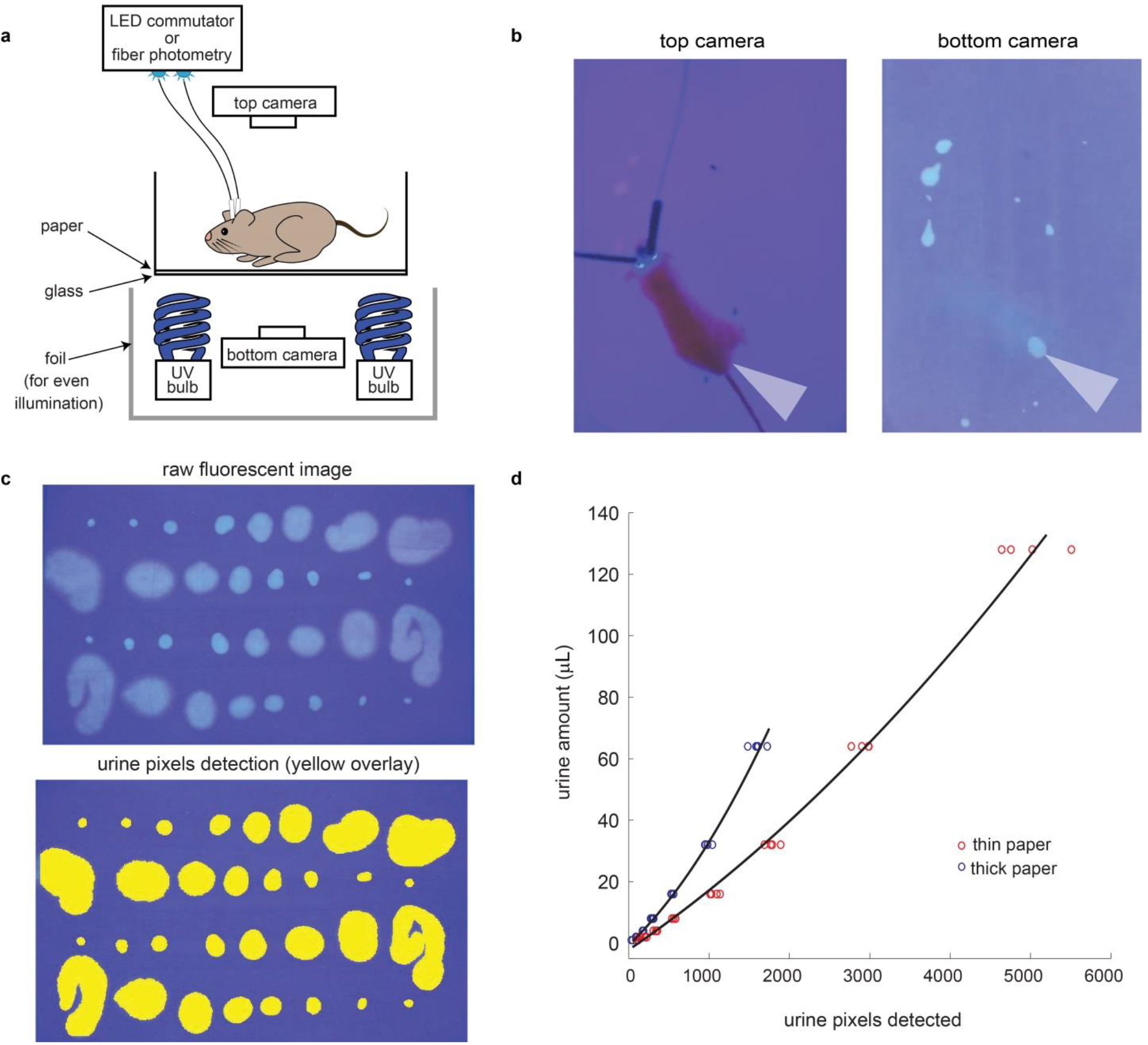
Visualizing and quantifying urination behavior. **a,** Schematic of behavioral setup for simultaneous optogenetics / fiber photometry, video recording, and analysis of urine excretion of awake behaving mice. **b,** Left: top camera records mouse position, right: urine fluoresces under UV light enabling excretion to be visualized throughout assay. Grey carrot indicates position between synchronized images from top and bottom cameras. **c,** Example of automated urine pixel detection with calibration data consisting of 4 replicates of 8 different volumes of male mouse urine on thin chromatography paper. **d,** Second order polynomial fit to calibration data on thick and thin paper; coefficients from these were used to calculate all urine amounts reported in microliters.

**Extended Data Fig 4.**
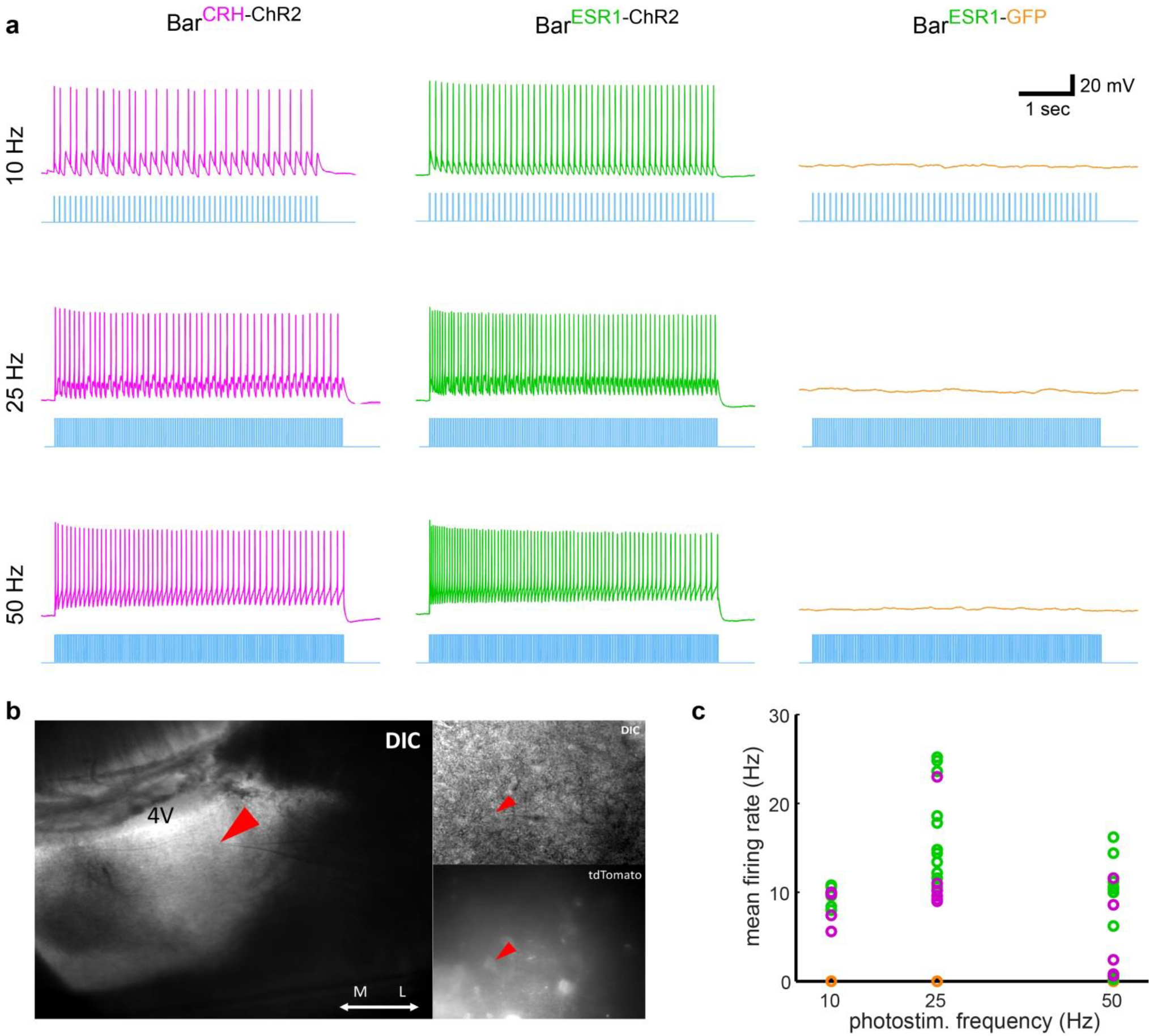
Whole-cell recordings of Bar^CRH-ChR2^ and Bar^ESR1-ChR2^ neurons during photostimulation. **a,** Example current clamp traces from representative Bar^CRH-ChR2^ (magenta), Bar^ESR1-ChR2^ (green), and Bar^ESR1-GFP^ (orange) neurons during 5 sec photostimulation bouts at 10/25/50 Hz. **b,** Visualization of recording location of Bar^ESR1-ChR2^ neuron in (a) showing ChR2-tdTomato expression. **c,** Summary of stimulated firing rate versus photostimulation frequency for all recorded Bar^CRH-ChR2^ (magenta, N = 6), Bar^ESR1-ChR2^ (green, N = 12), and Bar^ESR1-GFP^ (orange, N = 4) neurons.

**Extended Data Fig 5.**
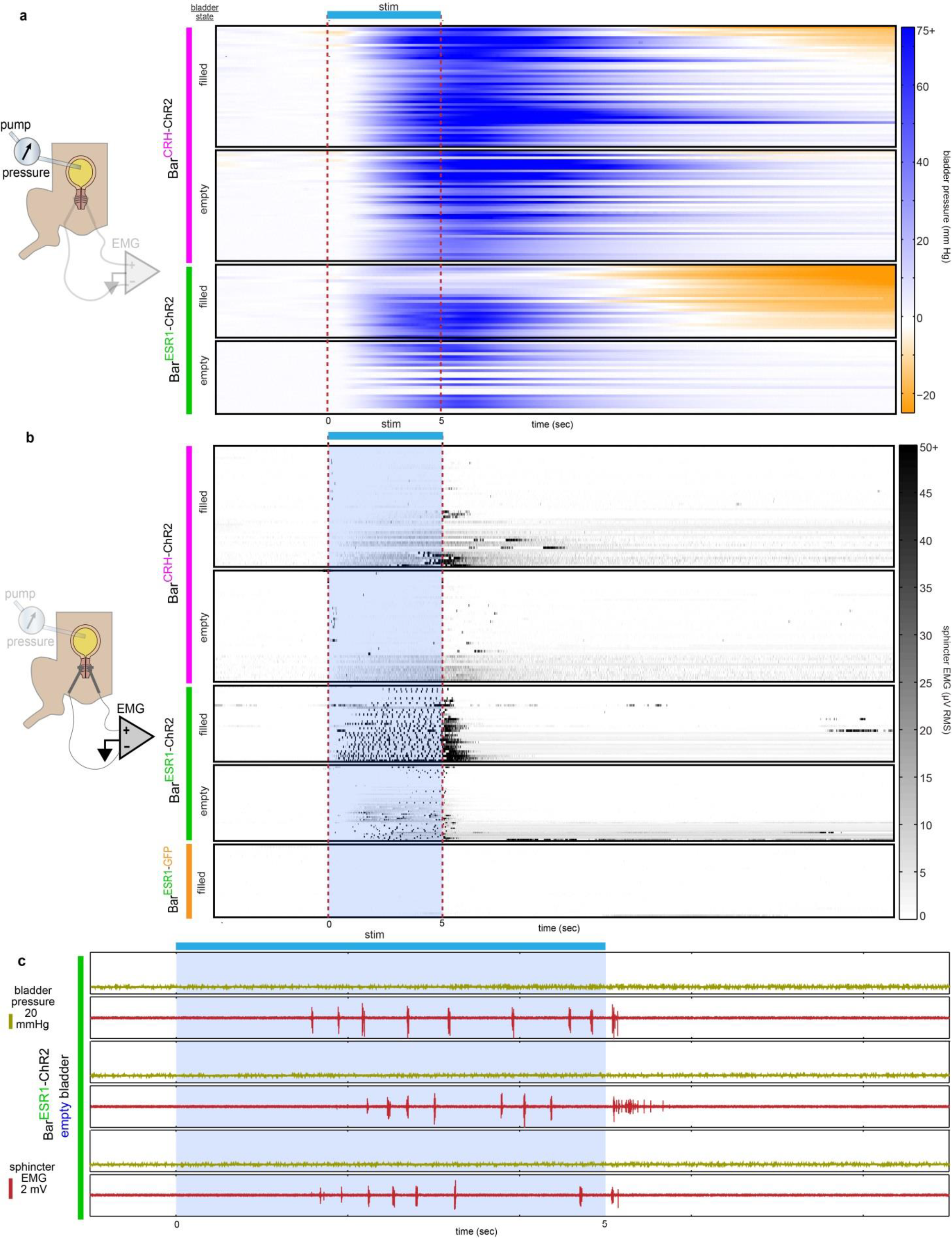
Bladder pressure and urethral sphincter EMG responses to Bar photostimulation. **a,** Schematic for optogenetic Bar stimulation during bladder pressure recording, and heatmaps of bladder pressure around photostimulation for all trials (N = 5 Bar^CRH-ChR2^, N = 3 Bar^ESR1-ChR2^ mice), in order of decreasing maximum pressure drop. Magenta bar on left denotes Bar^CRH-ChR2^ trials, and green bar denotes Bar^ESR1-ChR2^ trials, each with “filled” and “empty” bladder conditions (“filled” boxes are same data as Figure 3c). **b,** Schematic for optogenetic Bar stimulation during sphincter EMG recording, and heatmaps of sphincter RMS EMG around photostimulation for all trials (N = 5 Bar^CRH-ChR2^, N = 6 Bar^ESR1-ChR2^, N = 3 Bar^ESR1-GFP^ mice) in increasing mean voltage order. Magenta and green bars as in (a), and orange bar denotes Bar^ESR1-GFP^ control mice. **c,** Three example Bar^ESR1-ChR2^ photostimulation trials in the empty bladder condition, in which burst-like EMG activity was observed in the absence of bladder response. Top/yellow traces are bladder pressure, bottom/red traces are raw EMG. Blue shading and dotted lines delineate photostimulation periods.

**Extended Data Fig 6.**
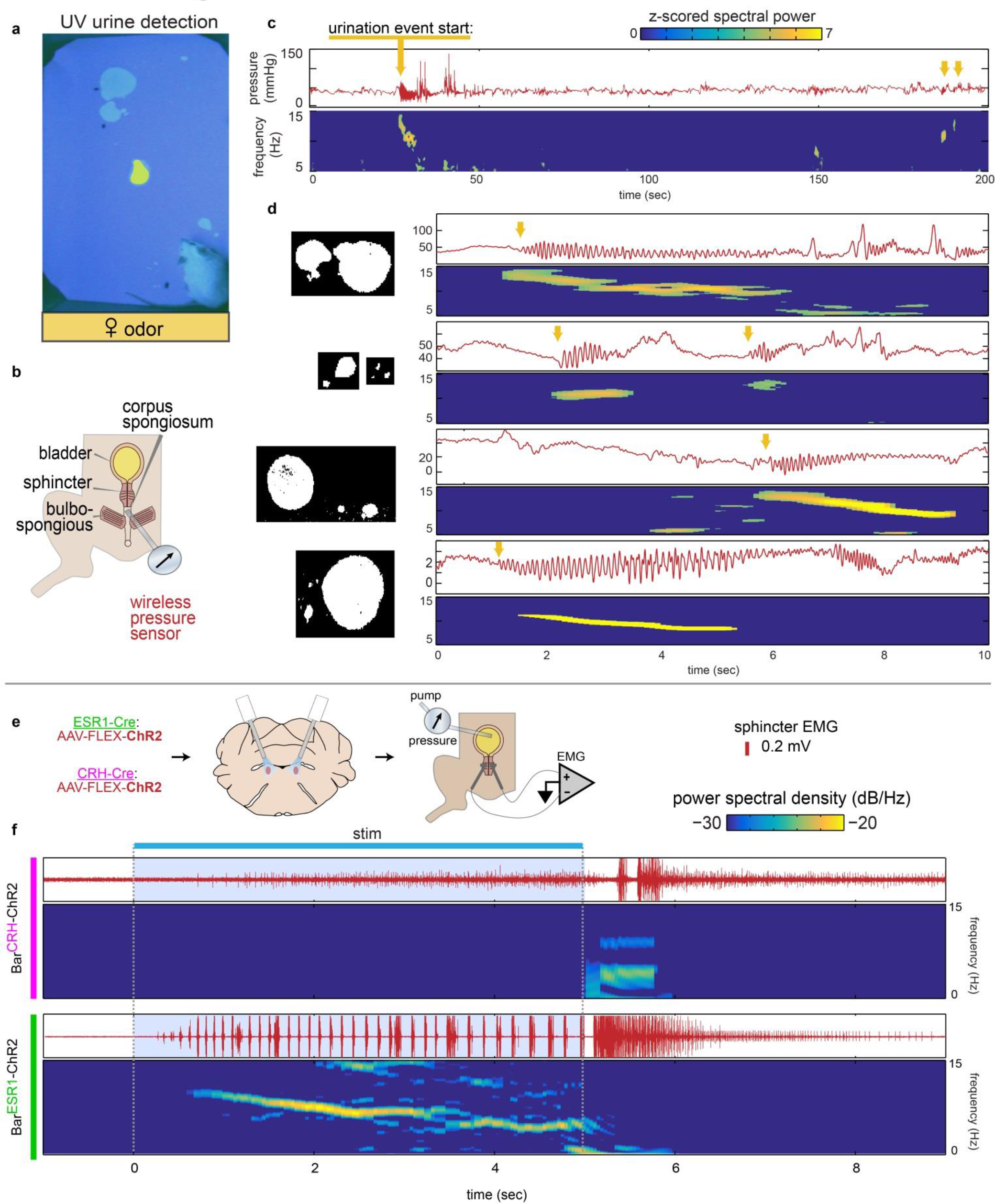
Frequency characteristics of urethral bursting during natural behavior and after Bar photostimulation. **a,** Example video frame from wireless corpus spongiosum pressure recording in the presence of female odor (yellow shading). **b,** Schematic of corpus spongiosum recording setup. **c,** Corpus spongiosum pressure recording after presentation of female odor. Top, raw pressure; bottom, spectral power in the 5-15 Hz band. Yellow arrows mark approximate start times for urination events. **d,** Shorter timescale recordings as in (c), for 5 urination events across 2 mice. Binary images on the left show relative sizes of thresholded urine marks corresponding to bursts on the right. Frequency typically decreases over a large burst lasting a few seconds, although shorter bursts were also observed (2^nd^ from top). **e,** Schematic for optogenetic Bar stimulation during sphincter EMG recording. **f,** Example raw sphincter EMG and corresponding spectral power in the 5-15 Hz band for photostimulated burst responses in Bar^CRH-ChR2^ (top) and Bar^ESR1-ChR2^ (bottom) mice. Bar^CRH-ChR2^ burst is preceded by an increase in tonic activity during the photostimulation period, whereas Bar^ESR1-ChR2^ burst occurs at low latency without preceding tonic activity, and displays decreasing frequency characteristic of natural bursts.

**Extended Data Fig 7.**
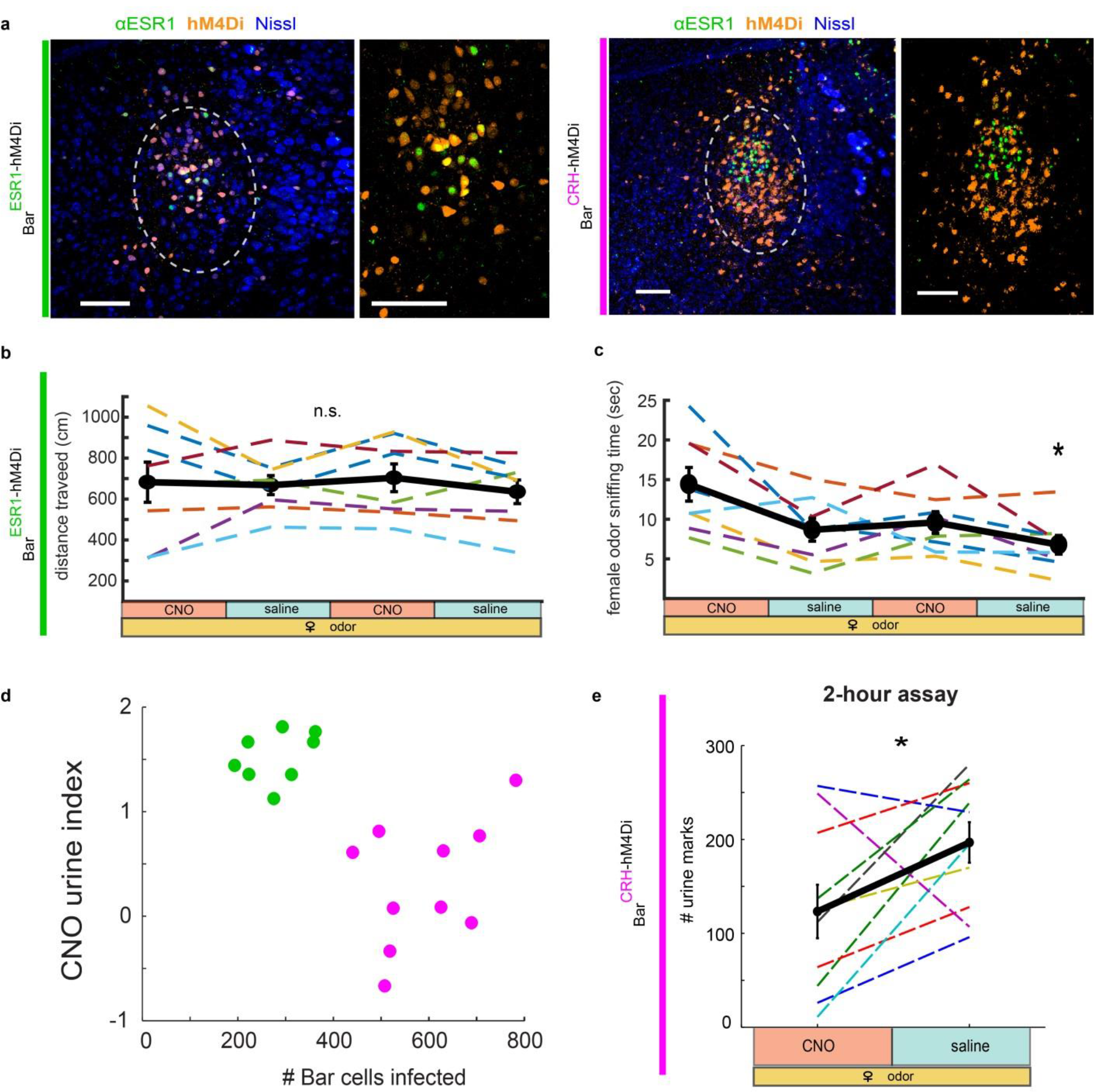
Viral infection and behavioral controls for Bar^ESR1-hM4Di^ and Bar^CRH-hM4Di^ chemogenetic inhibition. **a,** Example hM4Di expression in Bar of ESR1-Cre, left, and CRH-Cre, right, mice; larger views minus Nissl on the right of each full image. **b,** Total distance traveled during the assay shown in Figure 4e for Bar^ESR1-hM4Di^ mice (N = 8). Thin dotted lines are individuals, thick black line is mean ± s.e.m. **c,** Same as (b), but for total female urine odor sniffing time. **d,** Number of Bar cells infected with hM4Di virus versus CNO urine inhibition index (see Methods) for all mice (green for ESR1-Cre, magenta for CRH-Cre). **e,** Analysis of Bar^CRH-hM4Di^ mice (N = 10) injected with either saline or CNO on consecutive days prior to a 2-hour urination assay, similar to that previously published^10^ and which is not limited to odor-evoked, voluntary urination. n.s. = not significant, *p<0.05, n.s. p>0,05 (Friedman’s with Dunn-Sidak posthoc in panels b & c, Wilcoxon rank sum in panel e) for difference between CNO and saline treatment days. Scale bars = 100 μm.

**Extended Data Fig 8.**
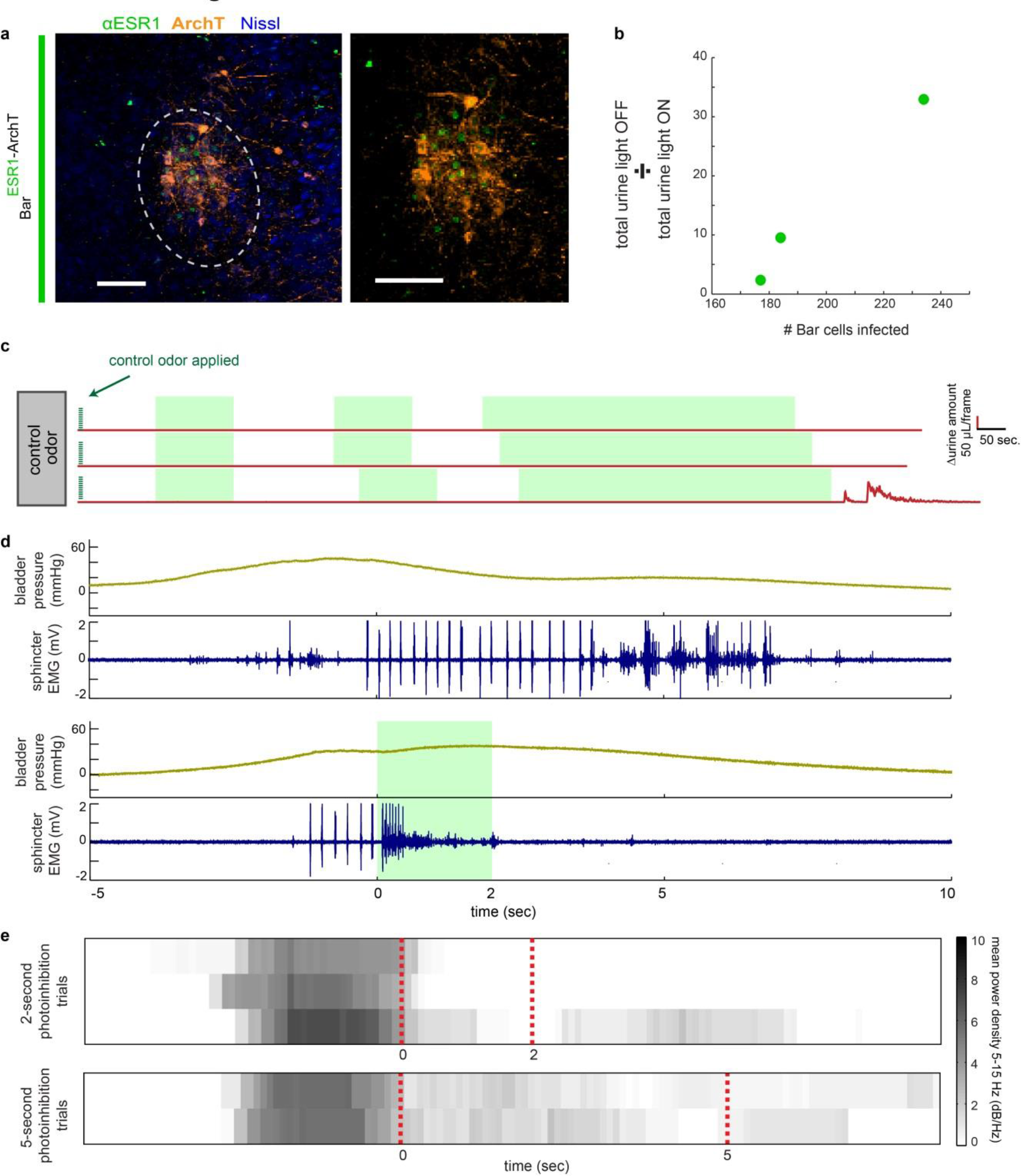
Bar^ESR1-ArchT^ photoinhibition terminates sphincter bursting during cystometry and does not result in rebound urination in awake mice. **a,** Example ArchT-GFP expression in Bar of ESR1-Cre mouse; right, larger views minus Nissl. **b,** Number of Bar cells infected with ArchT virus versus total urine with light OFF (not inhibited) divided by that with light on (while inhibited) for all mice (N = 3). **c,** Δurine amount around two 30 second photoinhibition periods followed by one 2 min. photoinhibition period, during which control odor only was present. N = 9 total photoinhibition bouts from 3 mice. **d,** Bladder pressure and sphincter EMG during cystometry, top, with natural unimpeded cycling, and bottom, in which 2 seconds of Bar^ESR1-ArchT^ photoinhibition was triggered as soon as filling evoked bursting was detected, which terminates bursting within ~100ms. **e,** Heatmap of mean EMG power density at bursting frequencies (5-15 Hz) during Bar^ESR1-ArchT^ photoinhibition as in bottom of panel d (top: 2 second inhibition trials; bottom: 5 second inhibition trials; N = 2 mice). Green shading or red dotted lines mark photoinhibition periods. Scale bars = 100 μm.

**Extended Data Fig 9.**
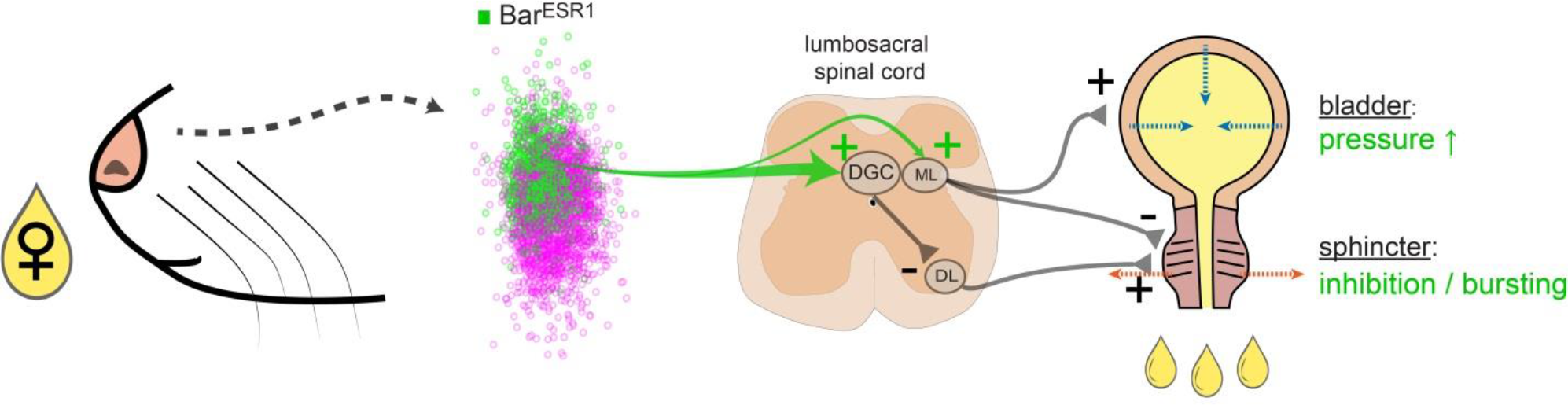
Simplified summary of a nose-to-sphincter circuit. Bar^ESR1^ (green) and Bar^CRH^ (magenta) neurons are intermingled, and the minority Bar^ESR1^ population projects to both the mediolateral column (ML) and heavier to the dorsal grey commissure (DGC), which directly inhibits sphincter motoneurons in the dorsolateral nucleus (DL). Activation of Bar^ESR1^ neurons increases bladder pressure and simultaneously inhibits the sphincter via bursting, thus driving efficient urine excretion. While it is likely that functionally distinct neurons in Bar act together in pelvic floor coordination, the minority Bar^ESR1^ population has the largest effect on urine output through its influence on the urethral sphincter. Heterogeneity in Bar may allow for more complex patterns of pelvic ganglia and pelvic floor muscle coordination during behaviors such as defecation, sex, and parturition. Thoracolumbar projections of Bar^ESR1^ neurons are not shown, as well as afferent feedback connections from bladder and urethra.

## Methods

### Animals

All animal procedures were conducted in accordance with institutional guidelines and protocols approved by the Institutional Animal Care and Use Committee at The Scripps Research Institute. BALB/cByJ male mice were group housed at weaning, single housed at 8 weeks old for at least 1 week before any testing, and maintained on a 12/12hr light/dark cycle with food and water available ad libitum. All mouse lines are available at The Jackson Laboratory: CRH-Cre (stock #: 012704,), Esr1-Cre^20^ (stock #: 017911), Vgat-Cre (stock #: 016962), Vglut2-Cre (stock #: 016963), ROSA-LSL-tdTomato (Ai9, stock #: 007909), and ROSA-LSL-ZsGreen (Ai6, stock #: 007906). CRH-Cre and ESR1-Cre mice were backcrossed into the BALB/cByJ background for 3+ generations.

### General surgical procedures

Mice were anesthetized with isoflurane (5% induction, 1-2% maintenance, Kent Scientific SomnoSuite) and placed in a stereotaxic frame (David Kopf Instruments Model 962). Ophthalmic ointment (Puralube) was applied, buprenorphine (Buprenex, 0.15mg/kg) was administered intramuscularly at the beginning of the procedure, and 500uL sterile saline containing carprofen (Rimadyl, 5mg/kg) and enrofloxacin (Baytril, 5mg/kg) was administered subcutaneously at the end of the procedure. Mice were monitored daily and given at least 14 days for recovery and viral expression before subsequent behavioral testing.

### AAV viral vectors

For photostimulation, AAV9-CAG-FLEX-ChR2-tdTomato (UPenn AV-9-18917P) was injected bilaterally at 1.4×10^12^ GC/mL in both ESR1-Cre and CRH-Cre animals. For CRH-Cre animals only, we also included AAV1-EF1α-FLEX-hChR2-eYFP (1:1 mix with above, UPenn AV-1-20298P) since this virus expressed at higher levels in Bar^CRH^ neurons in preliminary experiments. For photostimulation controls, AAV9-CAG-FLEX-GFP (UNC AV5220) was injected bilaterally at 3.2×10^13^ GC/mL in ESR1-Cre mice. For ESR1-Cre DREADD inhibition^34^, AAVdj-CAG-FLEX-hM4Di-GFP^35^ (Addgene plasmid # 52536, a gift from Scott Sternson) was produced by the Salk Institute Gene Transfer Targeting and Therapeutics Core (GT3) and injected bilaterally at 8×10^12^ GC/mL. We did not see efficient expression using this virus in CRH-Cre animals, so for CRH-Cre DREADD inhibition, AAVdj/1-EF1α-FLEX-hM4Di-mCherry (Addgene plasmid # 50461, a gift from Bryan Roth) was produced by Virovek and injected bilaterally at 4×10^12^ GC/mL. For photoinhibition, AAV9-CAG-FLEX-ArchT-GFP (UNC AV6222) was injected bilaterally at 2.2×10^12^ GC/mL in ESR1-Cre animals, and the same virus and titer were used for anatomical axon tracing unilaterally in both ESR1-Cre and CRH-Cre animals. For fiber photometry, AAV-CAG-FLEX-GCaMP6s^36^ (UPenn AV-9-PV2818) was unilaterally injected at 3.2×10^12^ GC/mL in ESR1-Cre animals.

### Viral injection and fiber optic implantation

Injections were made using pulled glass pipettes (tips broken for ID = 10-20 um) and a Picospritzer at 25 - 75 nL/min. For Bar injections, the overlying muscle was removed and a medial-lateral angle of 33° was used to avoid the 4^th^ ventricle. The pipette entry coordinate relative to bregma was 5.3mm caudal, 2.5mm lateral, and 3.2mm diagonally below the dura. The surrounding skull area was thinned for visualization with a diamond drill bit and the rostral-caudal coordinate was adjusted if necessary to coincide with the junction of the inferior colliculus and cerebellum, and to avoid hitting the transverse sinus. AAVs were injected 30-150nL per side, and the pipette was left in place for 5 min after injection, before slowly retracting. Fiber optic implants (4 mm length, Plexon 230 μm diameter for ChR2/ArchT and Doric 400 μm diameter for GCaMP) were inserted along the pipette track as above, 300 μm above the injection site for ChR2/ArchT, and 50μm for GCaMP. Additionally, two anchor screws (Antrin Miniature Specialties M1 X .060“) were attached over frontal cortex for animals with implants. After injection/implantation, the skull was covered with superglue and dental cement to seal the craniotomy and hold the implants in place.

### Spinal cord CTB injection

A 1-2 cm incision was made over lumbar segments, and the connective tissue and muscle overlying the vertebrae was minimally dissected^37^ to expose L1 and L2 vertebrae^38^. Vertebrae and underlying spinal segments were located by spinous process tendon attachments and spinous process shape, and confirmed by pilot injections of DiD dye. A spinal adapter^37^ for the stereotaxic frame (Stoelting 51690) was used to clamp L2 transverse processes, and a beveled glass pipette was lowered into the space between L1 and L2 vertebrae, 400 um lateral to the spinous process midline and 600 um below dura, to target the sacral mediolateral column bladder preganglionic neurons. After injection of 150 nL CTB-488 (ThermoFisher, 0.5% in PBS), the pipette was left in place for 5 min before slowly retracting, and then the injection site was covered with gelfoam and the overlying skin was sutured. Survival time was 5 days.

### Odor-motivated urination assay

Sexually naïve male mice were briefly prescreened for urination responses to 100uL female urine (>1 second odor sampling period with >3 urine marks within 1 minute) before any further testing or manipulation, which excluded 21% of all mice tested. The remaining 79% had surgical procedures and recovery or a 2 week waiting period before starting habituation. Mice were habituated in the behavior room for 3 consecutive days, for 16/8/4 minute durations on days 1/2/3. On day 3, control stimuli (100uL tonic water, which fluoresces under UV illumination) were pipetted from above at 0 min. and 2 min. and the baseline response was recorded. On subsequent test days, a 4 min. assay was used, with 100uL tonic water delivered at 0 minutes and 100uL female urine delivered at 2 min. All behavior was conducted during light hours under dim red light, and 70% ethanol was used to clean equipment between trials. The recording box consisted of a UV-opaque acryllic homecage with the bottom cut out, placed on top of 0.35mm chromatography paper (Fisher Scientific 05-714-4) resting on clear glass (Extended Data 4a). Two wide angle cameras (Logitech C930e), one above on a modified cage top, and one below the bottom glass, streamed video to a laptop computer at 15 frames per second, 640×360 pixel resolution. An analog pulse controlled LEDs in each camera field of view in order to synchronize cameras. Two UV fluorescent tube lights (American DJ Black-24BLB) surrounded by foil walls were used to evenly illuminate the chromatography paper from below. Videos were cut using Adobe After Effects and subsequently analyzed for urine marks using custom MATLAB software. The red and green channels of the RGB camera frames were used for urine detection, and the blue channel for mouse tracking. An output video with urine detection overlay was generated to manually verify automatic spot detection. Noldus Ethovision XT was used to automatically track mice and determine distance traveled and odor sniffing periods, defined as when the nosepoint occluded the female urine stimulus.

### Female urine collection

Adult (8-16 weeks) C57BL/6N female mice were housed 5 per cage, soiled male bedding was introduced into the cage 24 hours before the first collection night to induce estrous, and urine was pooled from 4 cages (20 mice total) over 4 days such that the stimulus consisted of a mix from all stages of the estrous cycle^39^. The mice were placed in metabolic cage for 12-16 hours at a time overnight, and urine was collected directly into a sterile tube on dry ice^39^ and temporarily stored at −20°C in the morning. After 4 consecutive nights of collection, urine was thawed on ice, rapidly passed through a 0.22um filter (Millipore Steriflip SCGP00525) before aliquoting and storing at −80°C. Two different batches of urine were collected for all experiments, and each was used with both control and experimental groups.

### Chemogenetic inhibition

After hM4Di^34^ viral injection, mice were allowed at least 21 days for recovery and expression, and then intraperitoneally injected 45-55 minutes before testing with either control saline plus 0.5% DMSO, or Clozapine N-oxide (CNO, 5mg/kg, Enzo Life Sciences BML-NS105-0025) in saline plus 0.5% DMSO. Control saline injections were performed on the 3 habituation days before female urine was given. Then on days 4/5/6/7, mice received CNO/saline/CNO/saline before the female urine countermarking assay described above. CRH-Cre mice were tested for 2 additional days (CNO, then saline) using the same assay but with 2-hour duration. Mice with less than 3 marks within 2 minutes after stimulus on both saline control days were excluded from analysis (8 of 34 mice), as well as mice that did not have bilateral hM4Di expression that spanned at least ±100um from the Bar rostral-caudal center, defined by ovoid Nissl clustering medial to locus coeruleus (7 of 34 mice). The “CNO Urine Index” (CUI) was calculated as [(% max. urine marks on saline days) - (% max. urine marks on CNO days)], such that CUI = 2 represents complete inhibition by CNO relative to saline, while CUI = 0 represents no difference between saline and CNO days.

### Optogenetic stimulation

For photostimulation experiments, fiber-implanted mice were briefly anesthetized with 5 % isoflurane before connecting and disconnecting patch cables (Plexon 0.5 m, 230 um diameter). An LED current source (Mightex BLS-SA02-US) driving two 465 nm PlexBright Compact LED Modules (Plexon) through a Dual LED Commutator (Plexon) provided 10±1 mW exiting the fiber tips. Optical power was measured (ThorLabs PM20A) before and after each session. Mice were placed in the same recording box described above for behavior, but with thinner 0.19 mm chromatography paper (Fisher Scientific 05-714-1). Initial experiments with different pulse widths determined 15 msec to be more effective than 5 msec or 1 msec at driving urination responses. All photostimulation bouts occurred for 5 sec duration using 15 msec pulses at five different frequencies: 1, 5, 10, 25, and 50Hz. These frequencies were stimulated in increasing order on the first day, and then repeated in decreasing order on the second day. At least 1 min elapsed between different photostimulation bouts, with additional delays occasionally necessary to allow the mouse to move to a clean section of paper. Videos were cut using Adobe After Effects and subsequently analyzed for urine marks using custom MATLAB software. Urine amount was calculated from urinated pixels detected using second-order polynomial coefficients determined with MATLAB polyfit on male urine calibration data (Extended Data 3c-d). Response latency was calculated as the earliest point when the normalized Δurine derivative reached 10% of maximum during the 15 sec response period. For a subset of mice, we repeated photostimulation on a third day under 1.5% maintenance isoflurane anesthesia. Four anesthetized 50 Hz/15 msec/5 sec photostimulation bouts separated by 1 min/1 min/1 min/5 min were conducted, then the isoflurane was removed and the mouse was allowed to recover to walking before waiting 5 min and following with two awake 50 Hz/15 msec/5 sec bouts separated by 1 min/5 min to confirm that awake urination was intact. After all experiments, mice were perfused and checked for viral expression and fiber placement as described for immunohistochemistry. Mice that did not have at least unilateral ChR2 expression that spanned ±100um from the Bar rostral-caudal center were excluded from analysis (9 of 29 mice).

### Optogenetic inhibition

For photoinhibition, all procedures were same as for photostimulation described above except for the following changes: fiber implanted mice were not anesthetized before connecting patch cables, but were habituated to the procedure for at least 3 days before testing. On the final habituation day, control odor was presented and 3 different photoinhibition periods were applied (2× 30 sec., 1× 2 min., separated by at least 30 seconds) to test the baseline effects of ArchT inhibition on urine output. Plexon 550 nm PlexBright Compact LED Modules were used, providing provided 6±1 mW exiting the fiber tips. During odor-motivated urination assay as described above, 2 min. of constant photoinhibition was applied 105 seconds after control odor, and 10-15 seconds before female urine. Urine marking behavior continued for 2 min. after photoinhibition ceased. Mice that did not have bilateral ArchT expression that spanned ±100um from the Bar rostral-caudal center were excluded from analysis (7 of 10 mice).

### Fiber photometry

Bulk GCaMP fluorescence was collected at 20 Hz using a similar setup to that previously described^40^. ΔF/F was calculated as (F - median(F) / median(F)) for each trial. An analog pulse controlled LEDs in each camera field of view as well as an Arduino sending triggers to the sCMOS camera (Hamamatsu ORCA-Flash4), in order to synchronize video and GCaMP data streams. Mice were recorded for 8 min. total (4 min. control odor only, then 4 min. with female urine stimulus). Δurine peaks were calculated from bottom video (MATLAB *findpeaks* function) with a minimum peak of 0.18 μL/frame, and GCaMP traces were analyzed around these peaks (zero lag) or at randomly selected times within the same assay (shuffle lag) as a control. The MATLAB *corrcoef* function was used to calculate correlation between GCaMP and Δurine traces.

### Electromyography and cystometry

Fiber-implanted mice were anesthetized with isoflurane (5% induction, 2% maintenance) and the bladder and external urethral sphincter (EUS, or urethral rhabdosphincter)^41,42^ were exposed via ~1cm midline abdominal incision. Flanged PE20 tubing connected to a syringe pump and pressure sensor (Biopac Systems DA100C/TSD104A) using a 25G needle was inserted and sutured into the bladder dome. Two tungsten wires (A-M Systems 795500) were stripped of insulation 1-2mm at the ends and inserted bilaterally (~2mm separation) into the EUS just proximal to the pubic symphysis, using a 30G needle. A third ground wire was stripped 3-4mm at the end and placed subcutaneously. The abdominal incision was sutured, allowing the tubing and wires to exit and connect to a differential amplifier (Biopac Systems EMG100C: gain = 5000, sample rate = 10kHz, low pass filter = 5kHz, 60Hz notch filter and 100Hz high pass filter). A digital input was simultaneously acquired at 10kHz, which was controlled by a TTL switch that also triggered optogenetic stimulation. After suturing, isoflurane was reduced to 1.0-1.8% (minimal to eliminate movement artifacts) and the bladder was filled at 10-20uL/min for at least 45 min. before starting photostimulation. Once a regular rhythm of urination cycles was established, the volume threshold was calculated as the mean volume of 3 cycles, and “filled” and “empty” states were defined as 75% and 10% of this mean value. Only mice with natural bursting cycles were analyzed for photostimulated or photoinhibited responses. Photostimulation consisted of 50 Hz/15 msec/5 sec photostimulation bouts separated by > 1 min. Photoinhibition consisted of constant illumination for 2 or 5 seconds, manually triggered at the beginning of a burst event. Root-mean-square (RMS) EMG traces were calculated using a 300 msec Gaussian filter and subtraction of the mean across 5 seconds prior to photostimulation. Sphincter relaxation periods were defined using RMS EMG data as periods between peaks >0.1mV (MATLAB *findpeaks* function) with amplitude less than the mean value prior to photostimulation. Frequency content of RMS EMG traces was calculated by first downsampling to 200 Hz, and then taking the FFT in overlapping 2 sec. rectangular windows. The spectrogram was thresholded at −40dB and burst duration was calculated as the time in which mean power in the 5-15Hz band is above this threshold.

### Wireless corpus spongiosum recording

Wireless pressure sensors (Data Sciences International, DSI PA-C10) were sterilized and implanted in the bulb of the corpus spongiosum that surrounds the urethra as previously described^43–45^, with the transmitter placed subcutaneously in the lateral abdominal area. After 1 week recovery, mice were recorded in the odor-motivated urine assay as described above, but with a single camera and UV illumination from above and the DSI RPC-1 receiver below the test cage. Pressure data was logged at 500 Hz and synchronized to urine imaging video. Frequency content of pressure traces was calculated by taking the FFT in overlapping 2 sec. hamming windows.

### Slice electrophysiology

Mice were deeply anesthetized with isoflurane, and acute 300μm coronal brain sections were prepared after intracardial perfusion of ice-cold choline-based slicing solution containing (in mM): 25 NaHCO_3_, 1.25 NaH_2_PO_4_, 2.5 KCl, 7 MgCl_2_, 25 glucose, 0.5 CaCl_2_, 110 choline chloride, 11.6 sodium ascorbate, 3.1 sodium pyruvate). Brains were quickly transferred and sliced in the same solution with a vibratome (LeicaVT1200).Sections were transferred to a recovery chamber and incubated for 15-20 min at 35 °C in recovery solution consisting of (in mM): 118 NaCl, 2.6 NaHCO_3_, 11 glucose, 15 HEPES,2.5 KCl, 1.25 NaH_2_PO_4_, 2 sodium pyruvate, 0.4 sodium ascorbate, 2 CaCl2, 1 MgCl2. Slices were maintained at room temperature for at least 30 min until transferred to bath for recording. Cutting solution, recovery solution, and ACSF were constantly bubbled with 95% O_2_/5% CO_2_. Slices were transferred to a recording chamber on an upright fluorescent microscope continuously perfused with oxygenated ACSF (in mM):125 NaCl, 25 NaHCO_3_, 2.5 KCl, 1.25 NaH_2_PO_4_, 11 glucose, 1.3 MgCl_2_ and 2.5 CaCl_2_ at 28-31 °C using a feedback temperature controller. Neurons labeled by fluorescent markers were visualized with a 40X water-immersion objective (Olympus) with epifluorescence and infrared differential interference contrast video microscopy. Recording pipettes were pulled from borosilicate glass (G150TF-4; Warner Instruments) with 3-5 MΩ resistance. The internal solution for current-clamp recording consisted of the following (in mM): 125 potassium D-gluconate, 4 NaCl, 10 HEPES, 0.5 EGTA, 20 KCl, 4 Mg_2_-ATP, 0.3 Na_3_-GTP, and 10 phosphocreatine. Recordings were made using a MultiClamp700B amplifier and pClamp software (Molecular Devices). The signal was low-pass filtered at 1 kHz and digitized at 10 kHz with a digitizer (Molecular Devices). For photostimulation of ChR2, 15 ms / 5 sec duration blue light pulses were emitted from a collimated light-emitting diode (473 nm; Thorlabs) driven by a T-Cube LED Driver (Thorlabs) under the control of a Digidata 1440A Data Acquisition System and pClamp software. Light was delivered through the reflected light fluorescence illuminator port and the 40X objective (light power at max setting measured at 13.45 mW). Analysis was performed in either Clampfit (Molecular Devices) or OriginPro 2016 (Origin Lab).

### Immunostaining

Animals were perfused with cold PBS followed by 4% PFA, and the brain / spinal cord (SC) was dissected and postfixed in 4% PFA at 4°C for 24-48 hours. The brain/SC was then washed in PBS and embedded in 1% low melting point agarose and cut on a vibratome at 50um for ESR1 and/or NeuN staining or 100um for Nissl-only staining. Spinal cords were cut transversely across the entire thoracolumbar and lumbosacral region and matched to segments using Nissl landmarks. For ESR1 immunostaining, free-floating sections were blocked in 1% BSA (Sigma A3059) in 1% PBST (PBS plus Triton X-100) for 3 hours, followed by primary incubation with anti-ESR1 antibody^17^ (antigen is mouse C-terminus fragment; Santa Cruz sc-542 or Lifespan C47042, rabbit polyclonal, 100ug/mL diluted 1:500 in 1% BSA / 0.3% PBST) overnight at room temperature. Sections were washed 3X with 0.1% PBST and blocked again at room temperature for 1 hour, before incubating in secondary antibody (ThermoFisher Alexa-Fluor 488 or 647 anti-rabbit IgG H+L diluted 1:2000 in 1% BSA / 0.3% PBST) at room temperature for 3 hours. Nissl stain (ThermoFisher NeuroTrace Blue or Deep Red diluted 1:200) was also included here if necessary, or incubated for 2 hours in 0.3% PBST if used alone. Sections were washed 2X in 0.1% PBST followed by 2X PBS, then mounted with ProLong Diamond (ThermoFisher). NeuN staining followed the same protocol as above but with NeuN primary antibody (EMD Millipore MAB377) diluted at 1:1000.

### Fluorescent in situ hybridization

Mice were anesthetized with isoflurane before rapid brain extraction, embedding in OCT, and freezing on dry ice. Coronal sections were cut at 20 um and stored at −80°C until processing according to the protocol provided in the RNAscope^®^ Multiplex Fluorescent v2 kit (Advanced Cell Diagnostics). Sections were fixed in 4% PFA, dehydrated, and hybridized with mixed probes: CRH (Mm-crh, Cat. 316091), Esr (Mm-Esr1-O2-C2, a 16ZZ probe targeting 1308-2125 of NM_007956.5.), Vgat (Mm-Slc32a1, Cat. 319191), and Vglut2 (Mm-Slc17a6-C2, Cat. 319171) for 2 h at 40°C and followed by amplification. Signal in each channel is developed using TSA Cyanine 3, fluorescein, and Cyanine 5 (PerkinElmer) individually. Sections were counterstained with DAPI and mounted with ProLong Diamond.

### Confocal Microscopy

Images were captured with Nikon A1 Confocal Microscope with a 10x air, 20x air or 40x oil objective. Nikon Elements software settings were optimized for each experiment to maximize signal range, and z-stack maximum projections were used for representative images and axonal projections while single optical slices were used for quantification of cell body overlap. For RNAScope, z-stacks were collected in 1 um increments throughout the z-axis.

### Anatomical quantification

The rostrocaudal center of Bar was defined as two consecutive 50 um section with greatest ESR1 and CRH-tdT labelling whenever possible, or by distinctive ovoid Nissl or NeuN boundaries. Custom MATLAB scripts were used to draw ROIs around Bar and semi-automatically count cells with clear cell body staining. Cells with high expression of ESR1 were distinguished from background labeling by thresholding in the ESR1 color channel just below the mean intensity level of nearby parabrachial neurons with established strong ESR1 expression^17,46^. Cartesian coordinates for cell locations were saved and the centroid of CRH-tdT cells was used to register different sections to generate the overlay plot in Fig. 1c and Extended Data 9. For calculation of fluorescence intensity ratio (Fig. 1k) in the lumbosacral mediolateral column (ML) and dorsal grey commissure (DGC), all intact L5-S2 sections with visible axons were used. A rectangular ROI was drawn using the Nissl color channel to encapsulate the MLs and area in between. This ROI was then equally divided into medial-lateral thirds and the Bar axon color channel was used to calculate the sum of pixel intensity across each third. The ratio was calculated as this total pixel intensity in the middle DGC third divided by that of the 2 ML thirds averaged together.

### Statistics and code availability

Nonparametric tests were used for all experiments. The Wilcoxon signed rank test (MATLAB *signrank*) was used for comparison of 2 paired groups, and the Wilcoxon rank sum test (MATLAB *ranksum*) for 2 unpaired groups. Friedman’s test (MATLAB *friedman*) was used to compare across CNO and saline treatments for 4-day DREADD experiments, followed by Dunn-Sidak posthoc tests (MATLAB *multcompare*). Points with error bars represent mean ± SEM. All analysis code is available upon request and will be deposited on GitHub.

## SI Guide

**Supplementary Video 1: Awake photostimulation of Bar^ESR1-ChR2^ neurons at five different frequencies.** This video shows a freely moving Bar^ESR1-ChR2^ mouse urinating in response to light pulses at five frequencies: 1,5,10,25,50 Hz. Photostimulation occurs when white frequency letters appear on middle-left. Top: recording from top camera (to visualize subject movement), Bottom: recording from bottom camera (to visualize and quantify urination). For schematic of assay chamber see Extended Data 3. Video condensed to 4X speed.

**Supplementary Video 2: Awake photostimulation of Bar^CRH-ChR2^ neurons at five different frequencies.** This video shows a freely moving Bar^CRH-ChR2^ mouse urinating in response to light pulses at five frequencies: 1,5,10,25,50 Hz. Photostimulation occurs when white frequency letters appear on middle-left. Video condensed to 4X speed in a two-camera recording set up as in Video 1.

**Supplementary Video 3: Anesthetized photostimulation of Bar^ESR1-ChR2^ neurons.** This video shows an anesthetized Bar^ESR1-ChR2^ mouse urinating in response to three 50Hz photostimulation bouts separated by 1 min intervals. The subject is moved between stimulations to observe and record urine excreted during each photoperiod. Video condensed to 4X speed in a two-camera recording set up as in Video 1.

**Supplementary Video 4: Photostimulation of Bar^ESR1-ChR2^ neurons during cystometry.** This video shows urine output during Bar^CRH-ChR2^ photostimulation while recording bladder pressure (top yellow rolling trace) and urethral sphincter EMG (bottom red rolling trace). Blue shading on plots and blue light shadow behind mouse delineate photostimulation period. Shorter sphincter bursting and urine excretion does not occur until after photostimulation. Video slowed to 0.67X speed show bursting clearly.

**Supplementary Video 5: Photostimulation of Bar^CRH-ChR2^ neurons during cystometry.** This video shows urine output during Bar^ESR1-ChR2^ photostimulation while recording bladder pressure (top yellow rolling trace) and urethral sphincter EMG (bottom red rolling trace). Blue shading on plots and blue light shadow behind mouse delineate photostimulation period.

**Supplementary Video 6: Odor motivated urination assay.** This video shows example urine marking behavior under UV light in response to control odor (1.5 - 2 min.) and female odor (2 - 4 min.) in a 4 min. assay. Video condensed to 4X speed in a two-camera recording set up as in Video 1.

**Supplementary Video 7: Photoinhibition of Bar^ESR1-ArchT^ neurons prevents rapid, odor-evoked urination.** This video shows a Bar^ESR1-ArchT^ mouse before, during, and after a 2 min. photoinhibition period during which female urine was presented. Video condensed to 4X speed in a two-camera recording set up as in Video 1.

